# Accounting for digestion enzyme bias in Casanovo

**DOI:** 10.1101/2024.05.16.594602

**Authors:** Carlo Melendez, Justin Sanders, Melih Yilmaz, Wout Bittremieux, Will Fondrie, Sewoong Oh, William Stafford Noble

## Abstract

A key parameter of any proteomics mass spectrometry experiment is the identity of the enzyme that is used to digest proteins in the sample into peptides. The Casanovo *de novo* sequencing model was trained using data that was generated with trypsin digestion; consequently, the model prefers to predict peptides that end with the amino acids “K” or “R.” This bias is desirable when the Casanovo is used to analyze data that was also generated using trypsin but can be problematic if the data was generated using some other digestion enzyme. In this work, we modify Casanovo to take as input the identify of the digestion enzyme, alongside each observed spectrum. We then train Casanovo with data generated using several different restriction enzymes, and we demonstrate that the resulting model successfully learns to capture enzyme-specific behavior. However, we find, surprisingly, that this new model does not yield a significant improvement in sequencing accuracy relative to a model trained without the enzyme information but using the same training set. This observation may have important implications for future attempts to make use of experimental metadata in *de novo* sequencing models.

## 1 Introduction

In tandem mass spectrometry (MS/MS) proteomics, a digestion enzyme cleaves proteins in a given biological sample into peptide sequences for further fragmentation and analysis. The enzyme trypsin is by far the most commonly used due to its reliability and stability, specificity in targeting lysine and arginine at the C-terminus of the peptide, and consequent tendency to yield peptides with a basic C-terminal residue which retains a positive charge, allowing for the generation of high quality spectra [1, 2].

Although trypsin is a standard protease for MS/MS, the use of alternative proteases and even cocktails of multiple proteases has been shown to improve peptide detection and proteome coverage in some settings [3, 4]. For example, the use of multiple digestion enzymes can allow for the creation of overlapping protein fragments, improving peptide detection [5]. Protein quantification has also been shown to improve using optimized mixtures of proteases beyond just tryptic digestion [6].

In general, for the task of peptide sequencing using database search, accounting for the digestion enzyme used in an MS/MS experiment is straightforward. Typical database search algorithms perform an *in silico* digestion of all the proteins in the database to generate a set of possible peptide sequences based on known digestion enzyme cleavage rules [2]. This approach works well, although it inevitably misses peptides that result from enzymes occurring naturally within the sample or from peptides created by non-standard enzymatic activity.

Explicitly accounting for these digestion rules is more challenging in the *de novo* sequencing setting, where a target peptide sequence is directly inferred from an observed mass spectrum. In recent years, deep learning models have become the preferred method for *de novo* sequencing [7–11], and these models are capable of implicitly learning the digestion rules associated with a given set of peptides from an MS/MS experiment. Although such models demonstrate excellent performance in sequencing proteins from MS/MS data, they are sensitive to the biases in their training data created by the use of a particular digestion enzyme.

Given the prevalence of trypsin as a digestion enzyme, most deep learning *de novo* sequencing methods have been nearly exclusively trained on tryptic data. That the models trained on this data learn a tryptic bias is sensible. For instance, given a spectrum that can be equally well explained by the sequences “PEPTIDEK” and “PEPTIDKE,” the former is *a priori* preferable given the digestion rules of trypsin. However, this preference for peptides that terminate with K or R will degrade sequencing performance in settings where the cleavage sites targeted by a digestion enzyme significantly differ from trypsin. In the example above, “PEPTIDKE” becomes the preferred sequence under digestion by gluC instead of trypsin. Therefore, the accuracy of these data-driven *de novo* sequencing models suffers when they are applied to non-tryptic data [12, 13].

The problem of augmenting deep learning *de novo* sequencing methods to account for the diverse digests generated by non-tryptic peptides has previously been handled in two different ways. First, one can train a single model on a training set containing a wide variety of digestion enzymes to try to account for as wide a range of digestion rules as possible. This approach has the advantage of potentially allowing the model to generalize to many different enzymes or enzyme combinations. Second, one can train a collection of models, with each one trained exclusively on data generated by a particular enzyme or enzyme cocktail. This approach avoids the need to balance different types of data during training; however, training a large collection of models is computationally expensive and may suffer from a lack of available data for less commonly used digestion enzymes. Moreover, this approach does not allow the model to generalize to settings where an MS/MS experiment makes use of novel combinations of digestion enzymes. Gueto-Tettay *et al*. experiment with both of these approaches, first training models on data generated using a single enzyme and then training on multi-enzyme datasets to increase their models’ generalizability [13].

In this work, we experiment with a third approach to account for enzymatic digestion: we provide the model with a representation of the identity of the enzyme used to generate each spectrum, and then we train the model to condition its predictions accordingly during training and inference. Specifically, we adapt the Casanovo *de novo* sequencing model to take into account the identity of the digestion enzyme used to generate a given precursor peptide. In the new model, Casanovo_*enz*_, each enzyme is mapped to a learned, high-dimensional latent representation, which is then passed as input to the model’s peptide decoder. To allow for data generated by cocktails of multiple enzymes, we sum the representations of each individual enzyme and pass the combined latent representation to the model’s decoder. We train Casanovo_*enz*_ on a dataset consisting of spectra generated from many different digestion enzymes and enzyme cocktails. We hypothesized that this modification of the Casanovo architecture would significantly improve generalizability to non-tryptic digests without degrading performance on tryptic data, outperforming a model that is trained from the same data but without the enzyme embedding.

Surprisingly, this hypothesis turns out to be false. Our empirical results suggest that providing Casanovo with explicit information about what digestion enzyme was used to generate a given spectrum yields only a very modest improvement in performance, relative to a model that is trained from the same data without any explicit enzyme information. We further provide evidence that the model successfully learns the digestion rules associated with a given enzyme, and that labeling spectra with the incorrect enzyme induces a predictable bias in the terminal amino acid distribution. Based on these results, we therefore trained a new version of Casanovo using data from a variety of different enyzmes, and we suggest that this single model be used for data generated using any enzyme or combination of enzymes.

Our motivation for reporting this negative result is three-fold. First, these results may help others to avoid carrying out similar experiments testing the same or closely related hypotheses. *A priori*, the idea of encoding various experimental parameters about a given spectrum, such as its associated collision energy or the precision of its *m/z* values, may seem like it would help boost *de novo* sequencing accuracy. But our results suggest otherwise. Second, and conversely, our results may spur others to come up with alternative, creative ways to represent and make use of such metadata in Casanovo or other *de novo* sequencing tools. Third, in the course of investigating the behavior of Casanovo and Casanovo_*enz*_, we uncovered evidence of significant batch effects, and these observations may be of more general interest. Finally, we also report on the training and public release of a new Casanovo model that performs substantially better than the previous model on data generated from non-tryptic digestion without any loss of accuracy on tryptic data.

## 2 Methods

### 2.1 Modifying Casanovo to account for digestion enzyme

Casanovo uses a transformer architecture to perform a sequence-to-sequence modeling task, translating from the sequence of peaks in a spectrum to the sequence of amino acids of the generating peptide. In Casanovo, each peak in an observed MS2 spectrum is treated as an element in a variable-length sequence. The *m/z* and intensity values of each peak are encoded into *n*-dimensional latent representations using, respectively, a collection of sinusoidal functions and a learned linear layer, and these encodings are summed. The encoded peaks are then input into the transformer encoder, where the transformer’s attention mechanism learns the context between pairs of peaks in the spectrum. The *n*-dimensional, contextualized peak encodings are then used as input to the transformer decoder for predicting the peptide sequence.

The decoding process proceeds in an iterative, autoregressive manner. We begin by providing the mass and charge of the observed precursor. Similar to the *m/z* and intensity values, the mass and charge are each encoded into *n*-dimensional latent representations using, respectively, a sinusoidal function and a learned linear layer, and these representations are summed. The transformer decoder uses the contextualized peak encodings and the precursor information to begin predicting amino acids of the peptide. Each amino acid is encoded into an *n*-dimensional latent representation by summing the outputs of two linear layers, one operating on a one-hot encoded representation of the amino acid and one operating on the amino acid position. To predict the amino acid at position *i*, the decoder takes as input *i n*-dimensional embeddings, representing the precursor mass and charge as well as the preceding *i −* 1 amino acids. Casanovo uses a beam search decoding strategy, where at each decoding step, we retain the top-scoring *k* beams, where *k* is a user-selected value. In each subsequent iteration, amino acids are added to the decoded peptide sequence, retaining the top *k* sequences until the decoded sequences for all of the beams have terminated or exceeded the precursor mass. Finally, the sequence with the highest score is retained as the putative peptide that generated the provided MS/MS spectrum.

To modify Casanovo to account for the identity of the digestion enzyme, we only need to change the decoding process. In the original model, the initial input to the decoder is the summed latent representations of mass and charge; in the extended model, we also include in this summation a latent representation of the enzyme. The enzyme identity is mapped to a *k*-dimensional latent vector via a learned embedding layer. For peptides digested by multiple enzymes we map each “component” enzyme to its own learned latent representation and then sum these latent vectors together, effectively treating the output as a superposition of the latent representations for each component. Everything else in the decoder, including the beam search procedure, remains unchanged. From the user’s perspective, this change requires specifying an additional “digest” option associated with a given spectrum file.

In this work, all of Casanovo’s hyperparameters (e.g., the number of transformers in the encoder and decoder, the number of heads per transformer, etc.) are as previously described [12]. The one exception is that, for efficiency reasons, the experiments reported in Sections 3.2–3.4 use six-layer rather than nine-layer transformers. When we train our final model in Section 3.5, we use the nine-layer architecture.

### 2.2 Augmenting Casanovo with an enzyme classifier

We implemented a variant of Casanovo that incorporates an enzyme classifier (Section 3.3). The classifier is a single, fully connected layer that takes as input the logits produced by the decoder transformer for the C-terminal amino acid. The classifier uses this information to predict which enzyme was responsible for generating the given spectrum. During training, this classifier is trained jointly with the rest of the Casanovo model by summing the two cross-entropy loss terms.

### 2.3 Data

In this work, we draw training, validation, and test data from two sources.

The first source, which we refer to as the “tryptic data,” is the data that was previously used in the development of Casanovo [12]. It consists of *∼*30 million PSMs from the MassIVE knowledge base (MassIVE-KB; v.2018-06-15) [14], all from experiments using trypsin. These 30 million PSMs were previously randomly split so that the training, validation and test sets are disjoint at the peptide level, yielding approximately 28 million training PSMs, 1 million validation PSMs, and 1 million test PSMs.

The second source, the “multi-enzyme data,” is a collection of *∼*1.1 million peptide-spectrum matches (PSMs) from MassIVE-KB v2.0.15, annotated as “proteomics experiments digested with various different enzymes.” The data includes PSMs from 812 mass spectrometry runs, using nine different digestion enzymes: arg-C, asp-N, chymotrypsin, elastase, glu-C, lys-C, lys-N, lys-arginase, and trypsin (Table 1). We first removed a set of 16,302 PSMs that had been produced by digestion with three different combinations of enzymes: arg-C and elastase; asp-N and lys-C; and glu-C and chymotrypsin. We then randomly split the remaining PSMs into training, validation, and testing sets in two stages. First, we placed PSMs involving peptides that appear in the MassIVE-KB data into the appropriate subset. Second, PSMs involving peptides that do not appear in the MassIVE-KB set were segregated into train, test and validation with a ratio of approximately 8:1:1. We used the multi-enzyme test set to evaluate tryptic bias in Figures 1 and Figure 5B, and we used the tryptic test set in Figure 5A. This procedure resulted in a train-test-validation split of, respectively, 883,281, 104,946, and 105,455 PSMs.

**Table 1:** Datasets employing various digestion enzymes. The columns indicate the total number of runs, the number of experiments (i.e., distinct MassIVE IDs), and the number of PSMs associated with each enzyme or combination of enzymes.

| Enzymes | Runs | MSV ID | PSMs |
| --- | --- | --- | --- |
| lys-C | 8 | 5 | 316873 |
| glu-C | 8 | 5 | 214657 |
| chymotrypsin | 8 | 3 | 180435 |
| arg-C | 8 | 2 | 114684 |
| lys-N | 6 | 1 | 49542 |
| asp-N | 8 | 2 | 45269 |
| lysarginase | 6 | 2 | 14967 |
| asp-N, trypsin | 6 | 1 | 10582 |
| glu-C, trypsin | 6 | 1 | 8863 |
| elastase, trypsin | 6 | 1 | 8027 |
| chymotrypsin, trypsin | 6 | 1 | 7923 |
| arg-C, asp-N | 6 | 1 | 7760 |
| asp-N, glu-C | 6 | 2 | 7745 |
| asp-N, chymotrypsin | 6 | 1 | 6377 |
| lys-C, elastase | 6 | 1 | 6326 |
| asp-N, elastase | 6 | 1 | 6276 |
| glu-C, lys-C | 6 | 1 | 6273 |
| asp-N, lys-C | 4 | 1 | 6233 |
| arg-C, glu-C | 6 | 1 | 6119 |
| glu-C, elastase | 6 | 1 | 5516 |
| elastase | 6 | 1 | 5488 |
| lys-C, chymotrypsin | 6 | 1 | 5437 |
| chymotrypsin, elastase | 6 | 1 | 5347 |
| arg-C, chymotrypsin | 6 | 1 | 5265 |
| glu-C, chymotrypsin | 4 | 1 | 5192 |
| arg-C, elastase | 4 | 1 | 4877 |
| arg-C, elastase, trypsin | 6 | 1 | 4346 |
| arg-C, lys-C, trypsin | 6 | 1 | 3917 |
| arg-C, chymotrypsin, trypsin | 6 | 1 | 3707 |
| glu-C, elastase, trypsin | 6 | 1 | 2726 |
| glu-C, lys-C, trypsin | 6 | 1 | 2658 |
| chymotrypsin, elastase, trypsin | 6 | 1 | 2416 |
| glu-C, chymotrypsin, trypsin | 6 | 1 | 2335 |
| arg-C, glu-C, lys-C | 6 | 1 | 2269 |
| arg-C, glu-C, trypsin | 6 | 1 | 2255 |
| glu-C, chymotrypsin, elastase | 6 | 1 | 2245 |
| lys-C, chymotrypsin, trypsin | 6 | 1 | 2245 |
| glu-C, lys-C, elastase | 6 | 1 | 2218 |
| arg-C, lys-C, elastase | 6 | 1 | 2072 |
| lys-C, chymotrypsin, elastase | 6 | 1 | 1906 |
| arg-C, chymotrypsin, elastase | 6 | 1 | 1854 |
| glu-C, lys-C, chymotrypsin | 6 | 1 | 1821 |
| arg-C, glu-C, chymotrypsin | 6 | 1 | 1808 |
| arg-C, glu-C, elastase | 6 | 1 | 1782 |
| lys-C, elastase, trypsin | 6 | 1 | 1721 |
| arg-C, lys-C, chymotrypsin | 6 | 1 | 1630 |
| Total |  |  | 1,109,984 |

**Figure 1.**
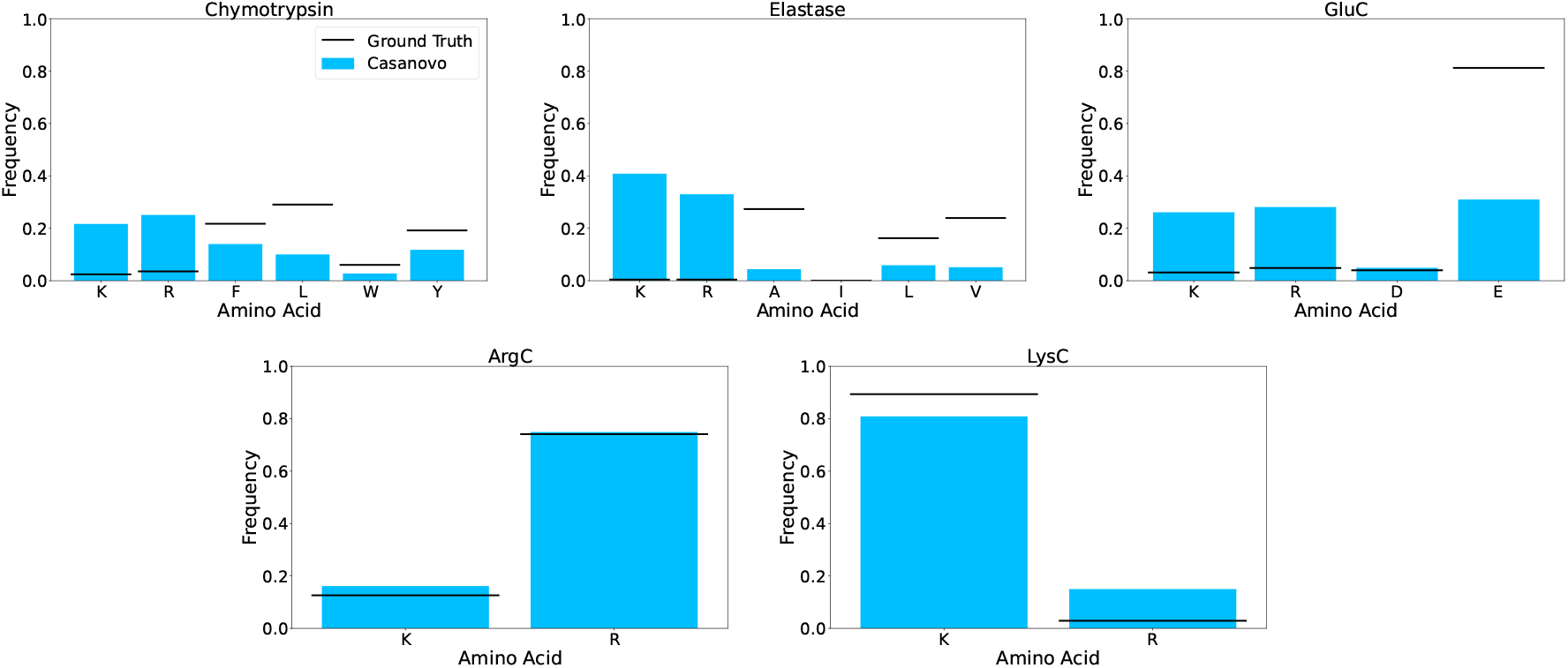
Casanovo has a tryptic bias at the C-terminus. Each each corresponds to a different digestion enzyme. Each vertical bar indicates the frequency of an amino acid at the C-terminus of Casanovo’s incorrect predictions. Each enzyme group includes bars for “K” and “R” as well as each of the expected C-terminal amino acids for that enyzme. Horizontal lines indicate the corresponding frequencies in the ground truth set of peptides.

For the final analysis in Figure 5, we used a simpler split of our original dataset, repeating the above procedure but without eliminating the 16,302 PSMs mentioned above. This resulted in a final train-test-validation split of 896,373, 106,933, and 106,678 PSMs, respectively.

For training and evaluation of Casanovo with an enzyme embedding (Figures 2–3, Section 3.2–3.3) we further downsampled our data to make trypsin less common. Starting from the combined multi-enzyme and trypsin training, validation, and test splits, we randomly downsampled the number of tryptic PSMs to 277,045 training and 27,437 test and validation set PSMs. These values were chosen to approximate those of the highest frequency enzyme classes (glu-C and lys-C) to prevent the problem of severe class imbalance due to tryptic training data. Finally, we created a test set of 200,000 PSMs from MSKB, subsampled from the aforementioned 1 million test PSMs, which we used for to evaluate the tryptic sequencing performance of all of our models.

**Figure 2.**
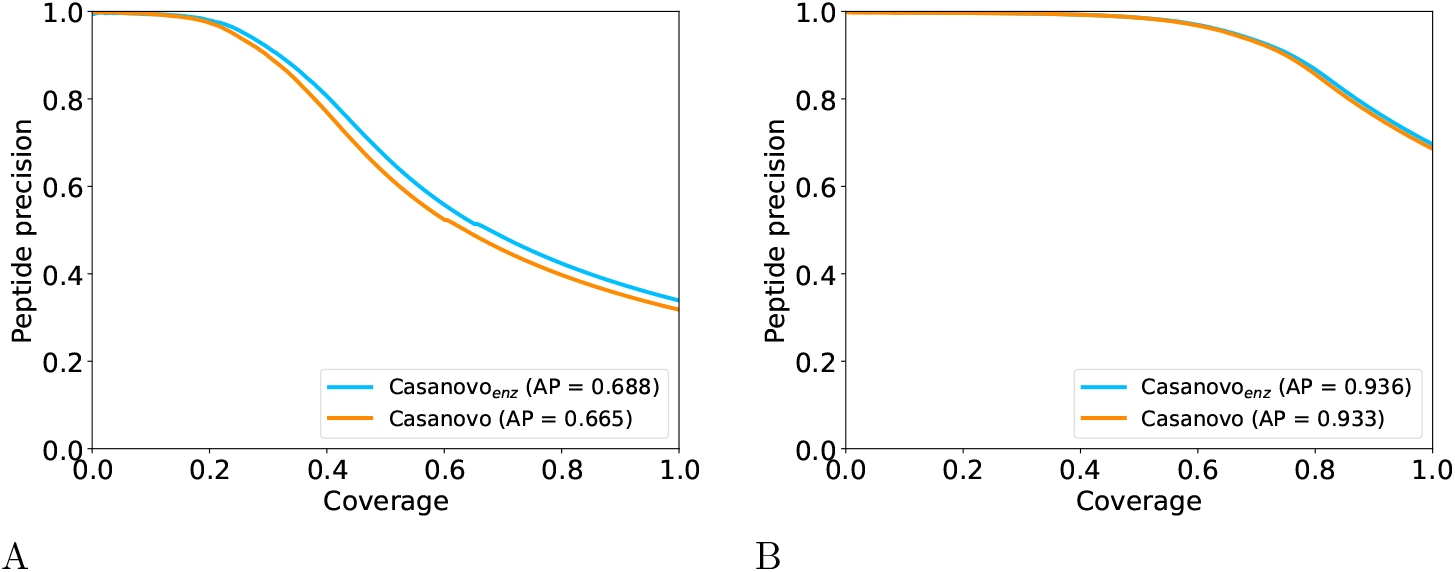
Comparison of Casanovo with and without the enzyme embedding. (A) The figure plots precision as a function of coverage, evaluated on the non-tryptic test set, for Casanovo and Casanovo_*enz*_. (B) Similar to panel A, but using the held out tryptic test set.

### 2.4 Batch-aware splitting procedure

In Section 3.4, to avoid leakage of batch-level information between the training and test set, we designed an additional train/test split that ensured that all spectra associated with a given MassIVE accession number were grouped together (Table 2). In this splitting procedure, we also aimed to keep all data associated with a given lab in either the train or the test set. Enzymes such as elastase and lysarginase were excluded entirely, because we only had data for these enymes from a single lab. For the remaining five enzymes, we designed the splits to allocate more spectra to the training than the test set. Ultimately, our training set contained 713,286 spectra, and our test set contained 120,658 spectra. This dataset was used to generate Figure 4.

**Table 2:** Details of the batch-aware splits.

| MassIVE ID | Enzyme | Spectra | PI | Subset |
| --- | --- | --- | --- | --- |
| MSV000083508 | arg-C | 110,664 | Kuster | Train |
| MSV000081607 | arg-C | 4,020 | Mirzaei | Test |
| MSV000083508 | asp-N | 26,126 | Kuster | Train |
| MSV000081607 | asp-N | 19,143 | Mirzaei | Train |
| MSV000081563 | chymotrypsin | 90,521 | Olsen | Train |
| MSV000083508 | chymotrypsin | 81,363 | Kuster | Train |
| MSV000081607 | chymotrypsin | 8,551 | Mirzaei | Test |
| MSV000083982 | glu-C | 8,015 | Xu | Train |
| MSV000086491 | glu-C | 63,169 | Xu | Train |
| MSV000081563 | glu-C | 90,576 | Olsen | Train |
| MSV000083508 | glu-C | 41,297 | Kuster | Test |
| MSV000081607 | glu-C | 11,600 | Mirzaei | Train |
| MSV000081563 | lys-C | 110,981 | Olsen | Train |
| MSV000083508 | lys-C | 63,016 | Kuster | Test |
| MSV000086491 | lys-C | 77,062 | Xu | Train |
| MSV000083982 | lys-C | 57,796 | Xu | Train |
| MSV000081607 | lys-C | 8,018 | Mirzaei | Train |

## 3 Results

### 3.1 The existing Casanovo model exhibits a tryptic bias

Before setting out to modify Casanovo to take in information about digestion enyzmes, we first quantified the extent to which the existing Casanovo model exhibits a tryptic bias. The model distributed with Casanovo version v4.1.0 was trained from a set of 28 million spectra, all of which were generated using tryptic digestion [12]. To evaluate the extent of enzymatic bias in this model, we ran Casanovo v4.1.0 on a data set consisting of 106,933 PSMs from MassIVE-KB v2.0.15, generated by five different enzymes. For this experiment, we excluded spectra generated using combinations of enzymes as well as enzymes with N-terminal digestion rules. For each enzyme, we considered only those PSMs for which Casanovo made an incorrect prediction, and we analyzed the frequency of different amino acids at the C-terminus of these predictions.

The results of this analysis suggest that Casanovo’s errors frequently involve incorrectly predicting a K or R at the C-terminus, in agreement with the tryptic digestion rule (Figure 1). Casanovo exhibits the greatest bias towards K and R and against the expected digestion sites for the enzymes chymotrypsin, elastase, and glu-C. For arg-C, which typically cleaves only after R, Casanovo over-predicts peptides that end in K. Conversely, for lys-C, which typically cleaves only after K, Casanovo under-predicts peptides that end in K and over-predicts peptides that end in R.

### 3.2 Adding enzyme embeddings to Casanovo does not improve performance on nontryptic data

We hypothesized that providing Casanovo with information about the digestion enzyme during training will allow the model to capture enzyme-specific biases in the distribution of terminal amino acids, improving the accuracy of predictions. To test this hypothesis, we trained a modified version of Casanovo, Casanovo_*enz*_, that includes the digestion enzyme embedding in the decoding step. The training set included the data sets described in Section 2.3: *∼*277,000 PSMs from MassIVE-KB and *∼*800,000 PSMs from the multi-enzyme training set. As a control, we also trained Casanovo on the same data, but without the use of an enzyme embedding. We then evaluated our two models, Casanovo_*enz*_ and Casanovo, on a held out multi-enzyme dataset consisting of 105,455 nontryptic PSMs generated with nine different digestion enzymes in 43 distinct combinations.

Suprisingly, the enzyme-aware model shows only a very small improvement in performance relative to the standard model trained on the same data (Figure 2A). In particular, we observe a 2.3% improvement in the average precision when we include enzyme information in the model. We also evaluated the performance of both models on a held out dataset of 200,000 PSM’s from MassIVE-KB generated with a tryptic digest. We find that the models yield nearly identical average precisions in this setting (Figure 2B).

To investigate whether including the enzyme embedding biases Casanovo_*enz*_’s predicted peptide sequences, we compared the distributions of C-terminal amino acids predicted by Casanovo and Casanovo_*enz*_ on the held-out test set. In this analysis, we segregated the spectra into groups based on the digestion enzyme and plotted the frequency of each C-terminal amino acid for the two models. The results of this experiment (Figure 3A–C) mirror the evaluations of the two models’ average precision on the multi-enzyme testset: both models perform approximately equally as well at predicting the expected terminal amino acids. To further investigate this enyzme-specific bias, we performed a second experiment in which we intentionally mislabeled some of the data at test time. Specifically, we extracted from the test set all spectra generated by glu-C, and we relabeled the data as chymotrypsin. The model’s C-terminal amino acid predictions show a marked shift towards the expected chymotrypsin distribution, which nominally cleaves after F, W, Y and L (Figure 3D). Thus, we conclude that the model is successfully learning the digestion rule associated with a given enzyme.

**Figure 3.**
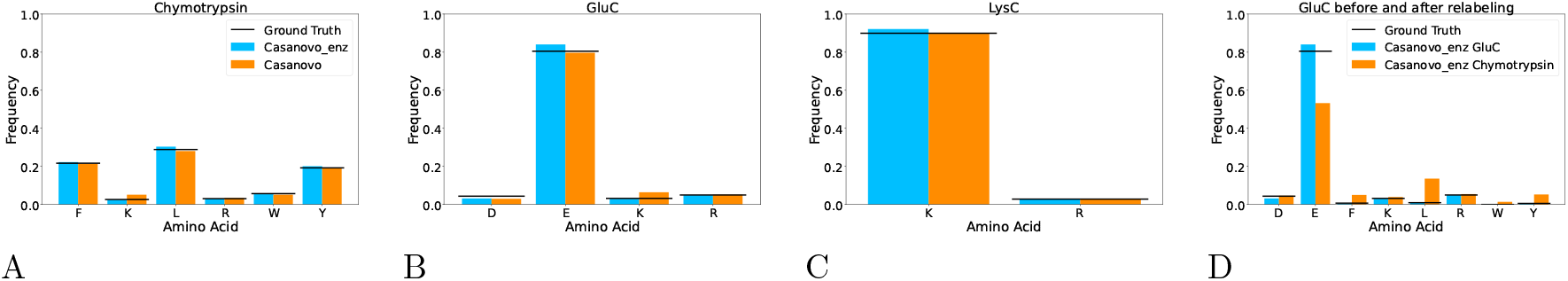
Evaluating C-terminal biases. (A–C) Each panel plots, for a given digestion enzyme, the frequency of C-terminal amino acids among the “true” peptides in the gold standard (horizontal line), the peptides predicted by Casanovo (orange), and the peptides predicted by Casanovo_*enz*_ (blue). Only the amino acids “K” and “R” plus amino acids corresponding to expected cleavage sites are included in each plot. The three panels correpond to the three enzymes with the most associated data. (D) Similar to panels A–C, but for glu-C data that was relabeled as being generated using chymotrypsin.

### 3.3 Modifying Casanovo to incorporate an enzyme classifier suggests that batch effects may be at play

One possible explanation for the relatively small performance improvement offered by the enzyme embeddings is that Casanovo_*enz*_ is not paying enough attention to the enzyme component of its input. We therefore designed a modified version of Casanovo with the aim of encouraging the model to retain and make use of the enzyme information. This modification involves adding to Casanovo a simple linear layer classifier that takes as input the latent embedding produced after decoding the first amino acid and aims to predict the identity of the enzyme associated with the spectrum (details in Section 2.2). The new model has two terms in its loss function: the original cross-entropy loss employed by Casanovo that compares the observed and predicted peptide sequences, plus a second cross-entropy term that compares the observed and predicted enzyme. We created two variants of this classifier-enhanced model, one that includes the enzyme embedding in its input (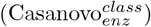) and one that does not (Casanovo^*class*^). Note that, because 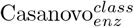 receives the enzyme identify as part of its input, we expected that the model should be able to easily re-identify the enzyme based on the latent embeddings produced during decoding.

We started by comparing the performance of the Casnovo_*enz*_ model with and without the auxiliary enzyme classifier. As expected, we found that the 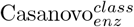 model does an excellent job at the enzyme classification task, achieving an accuracy of 100.0% when evaluated on an independent test set. However, we also observed that this same model performs slightly worse than a model that does not include the auxiliary classifier: the average precision on the test set changes from 68.8% for Casanovo_*enz*_ to 66.1% for 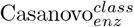. Thus, it appears that encouraging the model to retain the identify of the enzyme by incorporating the classifier in the loss function is not helpful.

Next we investigated the effects of adding the enzyme classifier to a Casanovo model that does not receive the enyzme identity in its input. We found that adding the classifier to the Casanovo model actually caused its performance to deteriorate by a substantial amount, with the test set average precision dropping from 66.5% for Casanovo to 50.4% for Casanovo^*class*^. Apparently, forcing the model to try to ascertain the identity of the associated enzyme, which is presumably a difficult task, hindered the model’s ability to carry out accurate *de novo* sequencing. Intriguingly, however, we also observed that Casanovo^*class*^ was surprisingly good at identifying the enzyme associated with a given spectrum, achieving a test set accuracy of 82.6%. Indeed, we conjectured that this accuracy is higher than should be achievable in principle, even in conjunction with a perfectly accurate *de novo* sequencer.

To test this conjecture, we carried out a simple experiment, in which we trained a classifier to predict the identify of the enzyme associated with a spectrum, when given as input the associated peptide sequence. Specifically, we encoded the peptide sequence into 60 features, with 20 representing the identity of each of the terminal amino acids, and 20 representing the counts of amino acids in the remaining (non-terminal) positions of the peptide. We used as a classifier a single linear layer similar to the one used in Casanovo^*class*^. When trained and evaluated on the same set of spectra, this classifier achieves an accuracy of only 58.2%, substantially lower than the 82.6% accuracy achieved by Casanovo^*class*^. From these observations, we conclude that when we ask Casanovo^*class*^ to identify the enzyme associated with a given spectrum, the model may instead be learning to identify batch effects associated with this spectrum.

### 3.4 After eliminating batch effects, the enzyme embedding is still not helpful

Based on these analyses, we returned to our original analysis, but we modified the way spectra are randomly partitioned with the aim of eliminating leakage of batch-level information between the training and test sets.

For this purpose, we used the MassIVE accession number as a proxy for batch effect, and we carried out a splitting procedure that places some MassIVE accessions in the training set and others in the test set while at the same time ensuring that (1) MassIVE accessions from the same lab are kept together, (2) each enzyme is represented in the training set and the test set, and (3) no peptides are shared between the training and test sets (details in Section 2.4). The splitting procedure placed five MassIVE accessions in the training set and two in the test set, yielding a total of *∼*713,286 training PSMs and *∼*120,658 test PSMs. The dataset contains data associated with the enzymes glu-C, arg-C, lys-C, lys-N, asp-N.

Repeating our training and testing procedure using Casanovo^*class*^ strongly supports the conclusion that, in our previous experiments, the classifier was learning to identify batch effects rather than inferring the enzyme identity: when we split in a batch-aware fashion, the classifier’s accuracy drops from 82.6% to 17.8%. It seems likely that the initial classifier was able to identify properties of the spectra that differ systematically across experiments.

In light of this result, we repeated the comparison of Casanovo and Casanovo_*enz*_. We reasoned that, now that the Casanovo cannot “cheat” by using batch-level information as a surrogate for the enzyme identity, the enzyme embedding might allow the Casanovo_*enz*_ to substantially improve its performance relative to the no-embedding model.

Surprisingly, this expectation turned out again to be incorrect: using the batch-aware splits, Casanovo_*enz*_ improves upon Casanovo by only a very small amount (1.2% average precision, Figure 4). The batch effects in our previous experiment led us to conjecture that Casanovo_*enz*_ was able to improperly infer the identity of the generating enzyme based on other properties of the spectrum, but having controlled for those batch effects, we now conclude that knowing the enzyme identity appears not to be as beneficial to Casanovo as we had expected *a priori*.

**Figure 4.**
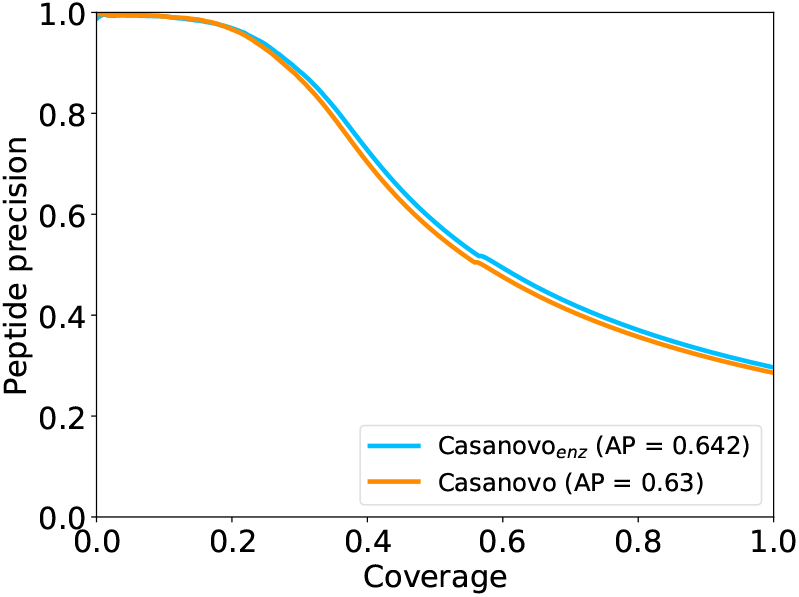
Comparing two Casanovo models using batch-aware splits.

### 3.5 Training a new version of Casanovo that includes multi-enzyme data

Providing a version of Casanovo that works well on data that was generated using enzymes other than trypsin is still an important goal. However, in light of the modest performance improvement that we observed from adding an enzyme embedding to Casanovo’s architecture, we opted for the simpler strategy of simply re-training Casanovo on training data generated using multiple enzymes. Specifically, we trained a standard 9-layer version of Casanovo on a combination of our multi-enzyme training set (896,373 PSMs from 46 different enzymes) and 2,000,000 tryptic PSMs randomly selected from MassIVE-KB. The training procedure converged after seven epochs of training. We compared the performance of our previous model, v4.0.0, and the newly trained model, v4.2.0, on our multi-enzyme test set as well as a test set of 200,000 MSKB PSMs.

The results of this experiment show that adding multi-enzyme data to the training set leads to no reduction in sequencing performance on tryptic data and markedly improved performance on non-tryptic data. On the tryptic test set, the two models exhibit very similar average precision values of 0.953 and 0.950 for v4.0.0 and v4.2.0, respectively (Figure 5A). In contrast, on the multi-enzymatic test set, the new model achieves a boost of 12.8% in average precision, from 0.561 for v4.0.0 and 0.689 for v4.2.0 (Figure 5B). Note that, because the peptide assignments for the tryptic and multi-enzyme data sets were made using different protocols, we do not expect the average precision values to be comparable between these two test sets.

**Figure 5.**
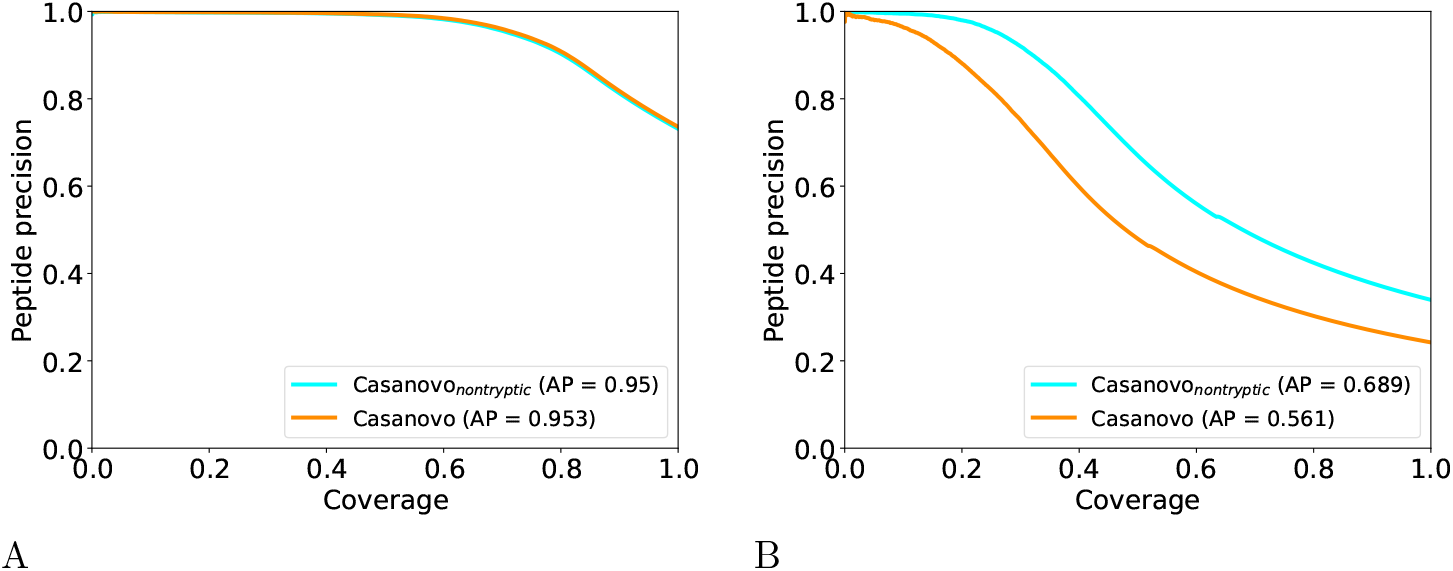
Performance of the old and new Casanovo models. (A) On a held-out test set of tryptic data, the old and new Casanovo models perform similarly. (B) On a test set containing non-tryptic data, the new model improves upon the old model.

## 4 Discussion

Our experiments lead us to several important conclusions. First, from a practical standpoint, it appears that providing Casanovo with information about the digestion enzyme provides only a very modest improvement in performance—on the range of 1-2% in our experiments. We made the decision not to push this modication into the released version of Casanovo because this small improvement must weighed against the additional complexity associated with adding this embedding functionality. Nonetheless, we provide a new model that delivers substantially improved performance on non-tryptic data while maintaining Casanovo’s excellent performance on tryptic data.

Several caveats come with this first observation. For example, the quality and variety of non-tryptic data could potentially impact the utility of this embedding; i.e., if we had 10 times the data of much higher quality then the embedding might be more useful. In addition, it is possible that hyperparameter optimization—selecting the dimensions of the model or the learning rate parameters—might have an impact on our results.

Our second conclusion is that the experiments in Section 3.2 suggest that Casanovo may be capable of identifying and exploiting batch effects. This observation has implications for how train/test splits are created for Casanovo and other deep learning *de novo* sequencing models, to ensure that test PSMs come from different experiments than training PSMs. This type of batch-aware splitting will likely be especially important for data-limited settings in which the data is derived from a small number of experiments. Thus, quantifying the impact of these batch effects on performance with respect to a larger variety of datasets seems like a potentially fruitful direction.

Although our primary result here is negative, we are not giving up entirely on the idea of encoding metadata into the input of Casanovo. Our experiments suggest that knowing the enzyme identity is not very helpful; however, other types of metadata—sample preparation details, liquid chromatography conditions, and instrument type and settings—may be useful in boosting the predictive accuracy of *de novo* sequencing methods like Casanovo. Exploring how best to incorporate such information into deep learning methods is a promising avenue for future research

## Acknowledgments

This work was funded in part by National Science Foundation award 2245300.

